# The multiple testing burden in sequencing-based disease studies of global populations

**DOI:** 10.1101/053264

**Authors:** Sara L. Pulit, Sera A.J. de With, Paul I.W. de Bakker

**Affiliations:** Brain Center Rudolf Magnus, Department of Neurology, University Medical Center Utrecht, The Netherlands; Brain Center Rudolf Magnus, Department of Psychiatry, University Medical Center Utrecht, The Netherlands; Center for Molecular Medicine, Department of Genetics, University Medical Center Utrecht, The Netherlands; Julius Centre for Health Sciences and Primary Care, Department of Epidemiology, University Medical Center Utrecht, The Netherlands

**Author notes:** Corresponding author: Sara L. Pulit; University Medical Center Utrecht Heidelberglaan 100 3584CX Utrecht The Netherlands *31 688 75 67745.

**Keywords:** *multiple test correction*, *association studies*, *complex traits*

## Abstract

Genome-wide association studies (GWAS) of common disease have been hugely successful in implicating loci that modify disease risk. The bulk of these associations have proven robust and reproducible, in part due to community adoption of statistical criteria for claiming significant genotype-phenotype associations. Currently, studies of common disease are rapidly shifting towards the use of sequencing technologies. As the cost of sequencing drops, assembling large samples in global populations is becoming increasingly feasible. Sequencing studies interrogate not only common variants, as was true for genotyping-based GWAS, but variation across the full allele frequency spectrum, yielding many more (independent) statistical tests. We sought to empirically determine genome-wide significance for various analysis scenarios. Using whole-genome sequence data, we simulated sequencing-based disease studies of varying sample size and ancestry. We determined that future sequencing efforts in *>*2,000 samples should practically employ a genome-wide significance threshold of of p <5 ×10^−9^, though the threshold does vary with ancestry. Studies of European or East Asian ancestry should set genome-wide significance at approximately p <5×10^−9^, but similar studies of African or South Asian samples should be more stringent (p <1×10^−9^). Because sequencing analysis brings with it many challenges (especially for rare variants), appropriate adoption of a revised multiple test correction will be crucial to avoid irreproducible claims of association.

## Introduction

In testing a single pre-specified hypothesis, researchers widely accept a p-value of *<*0.05 as sufficient evidence to reject the null hypothesis and make a claim of association. By testing an increasing number of hypotheses in a single experiment, however, she must account for the so-called testing burden of the experiment and accordingly adjust the significance threshold to reject the null hypothesis, thereby minimizing the chance of reporting a false positive (type I error). Though permutation is the ideal method for calculating an exact p-value, as it explicitly establishes the null hypothesis for a given test, this approach can be computationally expensive and time consuming. Thus, a widely accepted method to account for multiple comparisons (without permuting) is the Bonferroni correction, in which the experiment-wide p-value threshold is determined by dividing the desired type I error by the total number of independent tests performed.

Accounting for multiple testing has been a key issue in complex trait genetics over the last decade. Upon completion of the human genome map, candidate gene studies were widely employed to search for genes causing disease. Such studies involve genotyping single nucleotide polymorphisms (SNPs) in genes with a biologically plausible role in disease, and then testing for SNP-phenotype associations. However, in 2002, an extensive review indicated that only *~* 2% of candidate gene findings replicated (Hirschhorn et al., 2002), in part because of application of excessively liberal p-value thresholds and failure to account for testing many (potentially independent) SNPs. The development of genotyping arrays and efforts to catalogue commonly segregating SNPs (HapMap) in various global populations (The International HapMap 3 Consortium, 2010) allowed for broader interrogation of SNPs across the genome and in tens of thousands of samples through common variant genome-wide association studies (typically referred to as GWAS, though referred to here as common variant association studies, or CVAS (Zuk et al., 2014), for clarity). As CVAS developed, analysis groups used genome-wide significance threshold (p = 5 ×10^−8^) as a guideline for claiming a SNP as significantly associated to disease (Pe’er et al., 2008; Dudbridge and Gusnanto, 2008); this threshold reflects a Bonferroni correction for the approximately one million independent tests performed in a CVAS, holding type I error at 5%. Application of genome-wide significance has helped ensure that the majority of SNP-disease associations discovered through CVAS have been robust and reproducible.

The advent of sequencing technology has brought with it a new era in the study of human disease, and though sequencing remains more expensive than SNP array genotyping, dropping costs (Check Hayden, 2014) and new technologies (Loman and Watson, 2015) will soon allow for assembly of samples comparable to CVAS and study of populations mostly or entirely neglected by the field of complex trait genetics (i.e., non-European populations (Rosenberg et al., 2010; Pulit et al., 2010)). Through sequencing, disease studies are no longer limited to studying common variation (minor allele frequency (MAF) *>*5%) captured on SNP arrays but can now perform rare variant association studies (referred to here as RVAS (Zuk et al., 2014)) to test low-frequency (MAF 1 – 5%) and rare variation (MAF *<*1%) as well. Further, the data is no longer limited to single nucleotide variants (SNVs) but now also captures insertions and deletions (indels), copy number variants, and other structural variation. However, community standards for sequencing data analysis comparable to that of CVAS are still not in place, as RVAS can vary widely, from selection of sequencing platform, the interrogated genomic region, and depth of coverage (McKenna et al., 2010; DePristo et al., 2011; Van der Auwera et al., 2013). Because sequencing captures in principle the full frequency and variation spectrum (Marth et al., 2011), association testing in whole-genome data may include many more independent tests than are performed in CVAS. However, efforts to date seeking to determine the multiple testing burden in RVAS have focused exclusively on European-ancestry samples (Xu et al., 2014) or have relied on linkage disequilibrium pruning (Fadista et al., 2016), an imprecise measure compared to a permutation-based approach that explicitly calculates the null distribution in these studies.

Here, we will derive appropriate thresholds for whole-genome significance on the basis of simulated case-control studies of varying sample size and ancestry, using empirical whole-genome sequencing data collected in multiple populations and phenotypic permutations to establish the precise null distribution. The thresholds reflect an increased testing burden compared to CVAS, which have exclusively focused on common single- nucleotide polymorphisms.

## Methods

To simulate sequencing-based association studies with the aim of evaluating the multiple testing burden across various study designs (**Figure 1**), we used whole-genome sequencing data drawn from two studies representing five global populations (**Figure 2**).

**Figure 1:**
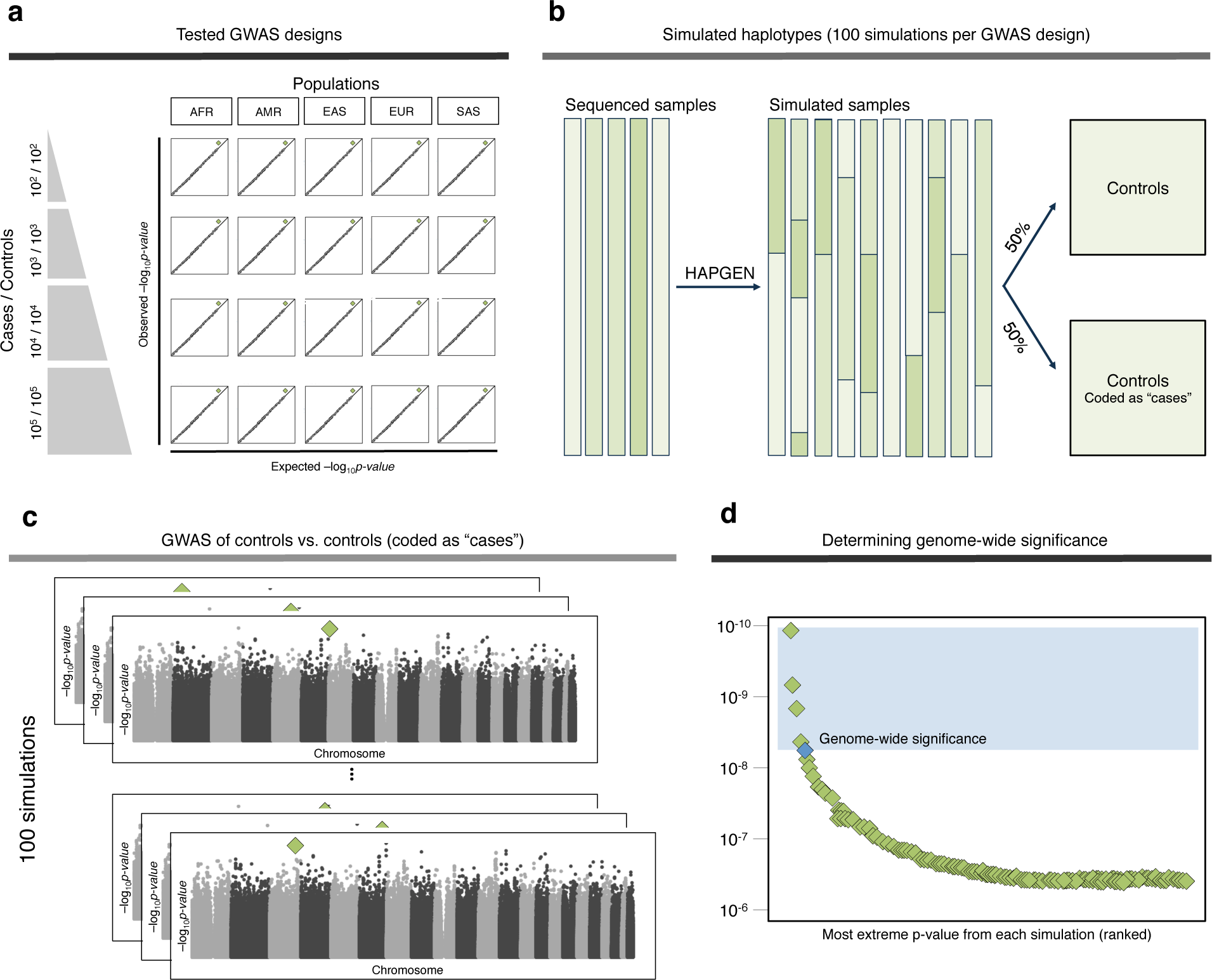
Pipeline to estimate genome-wide significance in sequencing-based rare variant association studies. (a) Twenty separate rare variant association study (RVAS) scenarios were simulated, varying the number of samples studied and the population (African ancestry, AFR; The Americas, AMR; East Asian ancestry, EAS; European ancestry, EUR; South Asian ancestry, SAS) analyzed. (b) Reference haplotypes from a single population were used to generate a group of controls that was then split into two equal subsets: one phenotypically coded as controls and the other phenotypically coded as cases (but still genotypically controls). (c) One hundred RVAS were run, comparing single nucleotide (SNV) frequency differences between controls and pseudo-cases. The most significant p-value (green diamond) was extracted from each of the 100 simulations performed for a given population- and sample-size specific study. (d) Each of these 100 p-values were then ranked; the fifth most significant p-value (blue diamond) represents genome-wide significance holding type I error at 5.

**Figure 2:**
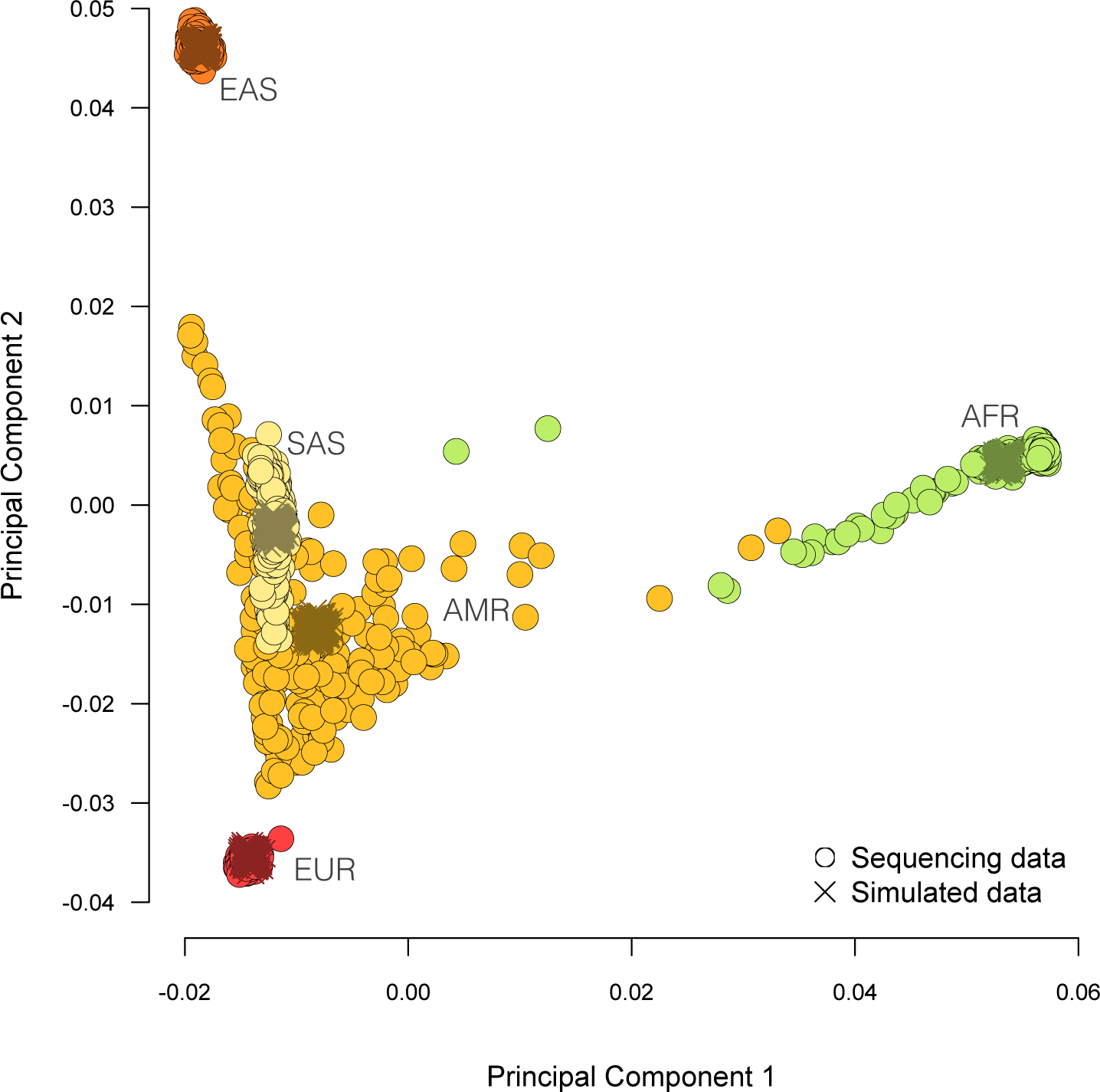
Principal component analysis of real and simulated data. Sequencing data from five global populations was used to simulate sequenced samples of different ancestries. We performed principal component analysis, projecting simulated data onto the real data, to ensure that global populations were being successfully simulated. AFR, African ancestry; AMR, The Americas; EAS, East Asian ancestry; EUR, European ancestry; SAS, South Asian ancestry.

To simulate sequencing studies focusing on European-ancestry (EUR) samples, we used data from the Genome of the Netherlands Project (GoNL; (Francioli et al., 2014)), which was comprised of 250 trios of Dutch ancestry whole-genome sequenced at *~* 14x coverage. To simulate sequencing efforts studying African-ancestry (AFR), samples from the Americas (AMR), East Asian ancestry samples (EAS), or South Asian ancestry (SAS) samples, we used data from the 1000 Genomes Project (1KG; (Auton et al., 2015)) Phase 3, which sequenced 2,504 unrelated individuals collected worldwide. Samples were sequenced at *~* 80x across the exome and *~* 4x outside the exome. Genotypes from both projects contained both SNVs and indels and had been phased (Menelaou and Marchini, 2013).

As the number of unrelated samples varied across the continental populations, we down-sampled each group by randomly selecting 268 unrelated individuals from each one, based on the smallest available continental sample (The Americas, N = 268). We then used HAPGEN (Su et al., 2011) to simulate various RVAS in each specific population at varying sample sizes (**Figure 1a**).

Using the whole-genome sequencing reference haplotypes and a recombination map, HAPGEN produced simulated whole genomes that maintained a similar frequency distribution and linkage disequilibrium (LD) structure to that observed in the reference. To enable explicit calculation of the empirical genome-wide significance threshold, we only simulated control samples. All controls were simulated together, and then split into two equal groups for genome-wide testing by randomly generating a binary trait that assigned half of the sample to a control group and assigned the other half as “cases” (but were still, genotypically, controls) (**Figure 1b**). We ran a principal component analysis (Price et al., 2006) on the real sequencing data, projecting sets of simulated samples on to the calculated principal components to ensure we were simulating samples reflective of the tested populations (**Figure 2**).

Once the simulations were complete and the phenotypes assigned, we ran a RVAS using PLINK (Purcell et al., 2007) across the autosomal chromosomes and chromosome X. A chi-square was used to test common variation (MAF ≥ 1%) and a Fisher’s test used to analyze rare variation (MAF *<*1%). For the GWAS comparing 100,000 controls and 100,000 controls, we used phenotypic permutations in PLINK1.9 (Chang et al., 2015) to evaluate the exact p-value at each variant with MAF *<*1%. Association tests were not corrected for any covariates. The final genome-wide p-value distribution was checked in initial simulations to ensure that it was uniformly distributed (**Figure 3**), as should be true under the null hypothesis of no association.

**Figure 3:**
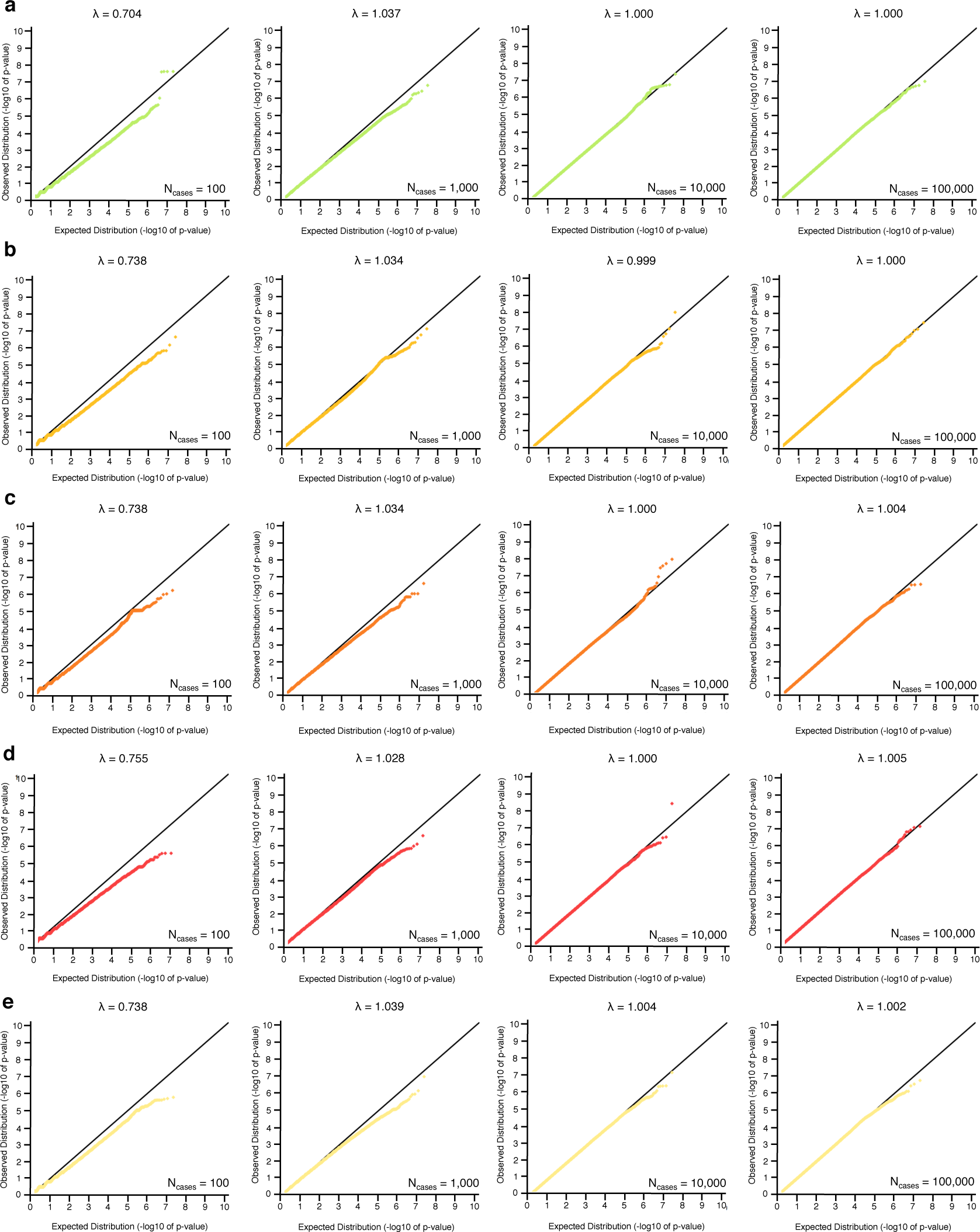
Quantile-quantile plots for a simulated RVAS within each continental population and selected sample size. The case-control ratio for each rare variant association study (RVAS) was held at 1:1. N_*cases*_ indicates the number of “case” (controls phenotypically coded as cases) included in the RVAS. (a) African-ancestry samples (AFR). (b) Samples from the Americas (AMR). (c) East Asian ancestry samples (EAS). (d) European-ancestry samples (EUR). (e) South Asian ancestry samples (SAS).

One hundred simulations were performed for each of the 20 scenarios tested (**Figure 1a**). To determine the genome-wide significance threshold for a single scenario, we first extracted the most extreme p-value from each of the 100 RVAS that had been performed for that specific sample ancestry and sample size. From these 100 p-values, we found the five most extreme p-values. The lower bound of these five p-values (i.e., the least significant of these five p-values) represents genome-wide significance for the specific study design when holding the type I error rate at 5% (**Figure 1c,d**).

## Results

Genome-wide significance for each of the 20 study designs tested is displayed in **Figure 4**. With the exception of studies using 1,000 controls vs. 1,000 controls, studies interrogating AFR samples always yielded the most stringent genome-wide significance threshold. This stringency is particularly evident in RVAS of 100,000 cases and 100,000 controls, where genome-wide significance is set for AFR samples at 1.237 ×10^−10^ and genome-wide significance is set for EUR samples approximately an order of magnitude lower, at 1.884 ×10^−9^. GWAS in EAS and SAS samples require quite similar genome-wide significance thresholds to that of EUR-based studies (**Figure 4**). Notably, for GWAS with 20,000 or 200,000 cases and controls, studies of AMR individuals required the second-most stringent significance threshold across all of the ancestry groups tested, potentially due to the non-European admixture in these samples.

**Figure 4:**
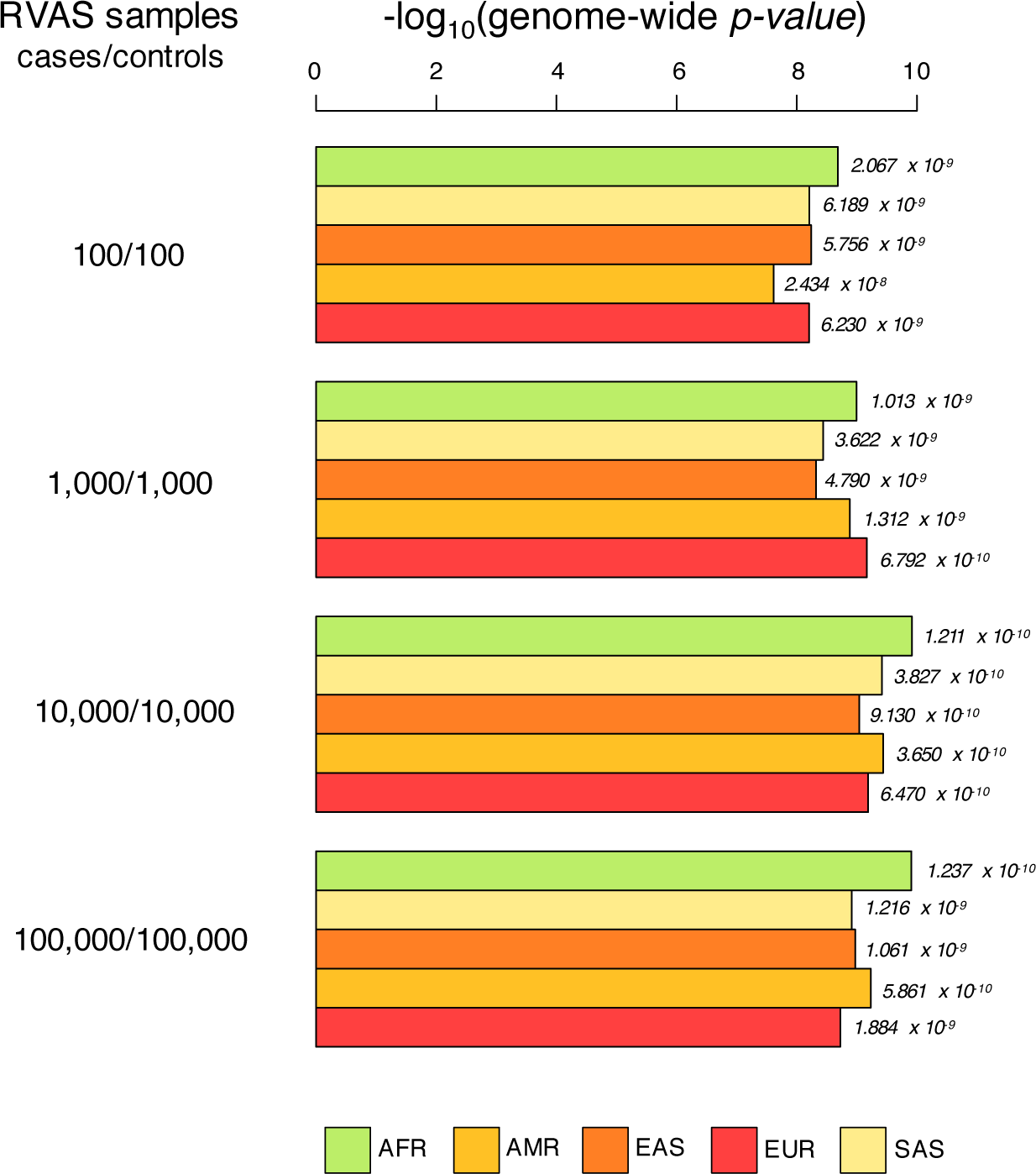
Genome-wide significance across five global populations in various rare variant association study settings. Rare variant association studies were simulated for various sample sizes and sample ancestries, maintaining a 1:1 case-control ratio. Genome-wide significance for the 20 simulated scenarios is shown. AFR, African ancestry; SAS, South Asian ancestry; EAS, East Asian ancestry; AMR, samples from the Americas; EUR, European ancestry.

The genome-wide significance threshold varied not only with study population but with sample size as well. Within each continental group, the genome-wide significance threshold became increasingly stringent as more samples were added to the analysis, up to the inclusion of 10,000 cases and 10,000 controls. Beginning from analysis of 100 controls vs. 100 controls and increasing to 10,000 cases and 10,000 controls, genome-wide significance increased by approximately an order of magnitude for each ancestry group studied. For example, studies of EUR samples successfully hold type I error at 5% by setting p = 6.230 *times*10^−9^ in 100 cases and 100 controls, but should adjust that threshold to p = 6.470 ×10^−10^ when testing 10,000 cases and 10,000 controls to control type I error. Interestingly, increasing from 10,000 cases and 10,000 controls to 100,000 cases and 100,000 controls had minimal impact on the whole-genome significance threshold, and even decreased its stringency (subtly) for a few of the ancestry groups tested (**Figure 4**). For example, genome-wide significance in 10,000 cases and controls of East Asian ancestry was p = 9.130 ×10^−10^ while it was effectively unchanged at p = 1.061 ×10^−9^ if 100,000 cases and 100,000 controls were tested. Similarly, the threshold for RVAS of the same sample sizes in AFR samples were p = 1.211 ×10^−10^ and p = 1.237 ×10^−10^, respectively.

## Discussion

Using whole-genome sequence data from two whole-genome sequencing projects, we simulated experiments in various populations to estimate the multiple testing burden in sequencing-based disease studies. Consistent with the population genetics observation that genetic diversity is higher in African-ancestry genomes (Campbell and Tishkoff, 2008), the testing burden in disease studies interrogating African-ancestry samples (or admixed samples with African ancestry) should be more stringent than in studies testing other ancestral groups. This increased burden reflects the increased number of independent variants in these genomes, also observed when determining the multiple testing burden in CVAS (Pe’er et al., 2008). Disease studies interrogating samples of South Asian ancestry should be similarly stringent. Large-scale sequencing studies, with samples numbering into the tens or hundreds of thousands and better powered to detect associations of small effect, should also consider using a genome-wide threshold stricter than the commonly used 5 ×10^−8^. This observation is also consistent with previous, similar analyses performed in common variant datasets (Pe’er et al., 2008), which note the relationship between effective sample size and the number of recombination events (and therefore, number of independent tests). RVAS with 20,000 cases and controls in our simulations were well-powered to achieve an essentially perfect null distribution (**Figure 3**). At such sufficient sample size, the genome-wide threshold appears to become robust to the addition of (potentially many more) samples.

Though HAPGEN can simulate genomes highly similar to genomes generated in a real sequencing project, using simulated rather than real genomes to determine multiple testing burden has its limitations. HAPGEN will only simulate variation similar to that contained in the reference haplotypes and will not inject new mutations into the data. Consequently, the larger simulated RVAS did not contain rare variation such as the large number of singletons and doubletons that would undoubtedly be detected in sequencing of many thousands of samples. Not testing these rare variants potentially underestimated the genome-wide significant threshold in the simulations containing *>*1,000 individuals, as they would be present and tested in a real study. Conversely, each RVAS tested all possible variants and not only common variation, likely deflating the estimate of genome-wide significance because of limited power in rare variant testing. Further, only 100 simulations were performed for each GWAS scenario, and more simulations would likely improve the precision of the estimates. The project differences between 1KG and GoNL may have also impacted the number of variants available for simulation and testing, though both projects had deep enough coverage to be sufficiently powered to detect the vast majority of variants with frequency *>*0.5% (Francioli et al., 2014; Abecasis et al., 2012; Auton et al., 2015).

Despite these drawbacks, we determined that for the GWAS in Europeans comparing 10,000 controls and 10,000 controls, the genome-wide significance estimate did not change after 1,000 simulations (data not shown). Our estimate for genome-wide significance in Europeans is also consistent with previously published work (Xu et al., 2014). Further, our method of simulating null GWAS by comparing controls versus controls yields more precise estimates of genome-wide significance than methods that rely on pruning SNPs in linkage disequilibrium (LD) to estimate the number of independent tests (Fadista et al., 2016). The number of estimated independent tests will vary significantly with the LD threshold selected to identify independent variants, and it is difficult to know which threshold will accurately identify all possible independent tests. Conversely, our approach relies on no such threshold selection, but rather directly reconstructs the null hypothesis at each tested variant in an RVAS. By recreating the null hypothesis, we need only select the type I error threshold (5%, as is typically done for RVAS and CVAS alike), to measure the multiple testing burden. The thresholds derived here are also penultimate thresholds and will likely be adjusted one last time, as detection of additional variation improves and we begin testing not only SNVs and indels that may be associated to disease (as was done here), but copy number variants, de novo mutations, repeats, and other more complex variation as well.

One analytic approach not addressed here is that future genome-wide significance thresholds may be informed by variant annotation. The thresholds presented here are set for the full set of single nucleotide variants (SNVs) tested, agnostic to function. Deriving function-specific thresholds, more nuanced than a single threshold because they are influenced by prior expectation of a variant’s biological impact, may improve power for discovery (Sveinbjornsson et al., 2016). We also do not address genic burden testing approaches, designed to test a collection of rare variants in the same gene or window in association to disease (Lee et al., 2014). Burden tests improve power by aggregating rare variation and reducing multiple testing burden by only testing the ~ 20,000 genes in the genome. A genome-wide significance estimate already exists for these tests (p ~ 4 ×10^−8^; (Xu et al., 2014)). Though burden tests improve power through variant aggregation, it remains difficult to know which variants exactly should be included in such a test. Functional annotation remains imprecise (Grimm et al., 2015), and incorrectly annotated variants (that are benign in actuality) will decrease power in a burden testing framework. Thus, with sufficient sample collection and sequencing, single-variant association testing will likely replace burden testing, as it allows for direct testing of causal variants, rather than association testing in collections of (possibly noisy) variants.

The genome-wide significance threshold has proved integral to the success of CVAS to date. In part because a SNP can only be claimed associated to a trait if it’s p-value surpasses 5 ×10^−8^, the majority of CVAS claims appear repeatedly at genome-wide significance as larger studies are performed (Ripke et al., 2014; Wood et al., 2014). As studies of common disease increasingly progress to genome-wide testing approaches that include both CVAS and RVAS, complex trait genetics has been invigorated by the promise that sequencing will reveal substantially more of disease’s genetic architecture. Yet sequencing brings with it many challenges. Statistical power to discover risk variants is limited (Goldstein et al., 2013; Kiezun et al., 2012; Kryukov et al., 2009), as risk variants will likely be of modest effect and low frequency (Steinthorsdottir et al., 2014; Sulem et al., 2011; Do et al., 2015). Further, the genome is filled with so-called “narrative potential” (MacArthur et al., 2014), mutations that have interesting biological consequence, such as nonsense mutations and frameshift indels, but that are phenotypically benign (MacArthur et al., 2012; Gratten et al., 2013) and even carried in randomly ascertained populations (Francioli et al., 2014; MacArthur et al., 2012). The challenges of sequencing, posed by both analysis and interpretation of the data, in addition to the mounting conversation to improve the reliability of published and publicly-available scientific research (Macilwain, 2012; Yong et al., 2013; Nature Neuroscience Editors, 2013; Pulit et al., 2014), make it crucial to have community consensus for claims of association made in sequencing studies in complex traits. Setting such a standard will lead to reporting of claims likely to replicate that are worthy of further study and may lead to novel insights into etiology, treatment, and prevention of disease.

